# Intrauterine exposure to hyperglycaemia in pregnancy and risk of adiposity in the offspring at 10 years of age – A community based retrospective cohort study in Sri Lanka

**DOI:** 10.1101/618991

**Authors:** Himali Herath, Rasika Herath, Rajitha Wickremasinghe

## Abstract

**Background:** Intrauterine exposure to a hyperglycemic environment can cause long term changes in body composition resulting in increased adiposity and cardio metabolic risk in the offspring. The aim of this study was to determine the association between hyperglycaemia in pregnancy (HIP) and risk of adiposity in the offspring at 10-11 years of age.

**Methods:** A retrospective cohort study was conducted in the Colombo district, Sri Lanka. 7205 children who were born in 2005 were identified through schools and Public Health Midwives in the community. Mothers of these children still possessing antenatal records were interviewed and relevant data were extracted from medical records to identify eligible participants. Exposure status (hyperglycaemia in pregnancy) was ascertained based on client held antenatal records. 159 children of mothers with HIP (exposed) and 253 children of mothers with no HIP (non-exposed) were recruited. Height, weight, waist circumference and triceps skin fold thickness (TSFT) of participants were measured to ascertain outcome status.

**Results:** The mean ages (SD) of exposed and non-exposed groups were 10.9 (0.3) and 10.8 (0.3) years respectively. The median BMI (17.6 vs 16.1, p<0.001), waist circumference (63cm vs 59.3 cm, p<0.001) and triceps skinfold thickness (13.7mm vs 11.2mm, p< 0.001) were significantly higher in the exposed group than in the non-exposed group.

Children who were exposed to intrauterine hyperglycaemia were more likely to be overweight (aOR=2.5, 95% CI 1.3-4.7), have abdominal obesity (aOR=2.9, 95% CI 1.2-6.8) and high TSFT > 70^th^ centile (aOR=2.1, 95% CI 1.2-3.9) at 10-11 years of age than children who were not exposed after adjusting for maternal BMI, birth weight and birth order.

**Conclusions:** Intrauterine exposure to HIP is associated with significantly higher risk of adiposity in the offspring at 10 years of age.

## Introduction

Hyperglycaemia in pregnancy (HIP) is one of the commonest medical conditions encountered in pregnancy. The International Diabetes Federation (IDF) estimates that one in six live births (16.2%) in the world and one in four live births (24%) in South East Asia are complicated with some form of hyperglycemia in pregnancy [1]. The majority (84% - 86%) of cases of hyperglycaemia in pregnancy is due to gestational diabetes mellitus (GDM) while the remaining cases are due to diabetes in pregnancy (DIP) which is either pre-existing type 1 or type 2 diabetes or diabetes first detected at any time during the index pregnancy [1,2]. The number of women having hyperglycaemia in pregnancy is increasing as a result of the increasing prevalence of obesity and diabetes in women and higher age at childbirth [3].

Pederson’s hyperglycemia-hyperinsulinism hypothesis, as proven by several studies, is still the basis of research on feto-maternal metabolism [4,5]. This hypothesis postulates that deficiency of maternal insulin causes a rise in maternal glucose, which in turn increases fetal glucose levels. This results in fetal hyperinsulinaemia which stimulates fetal growth and adiposity. Frienkel and Metzger stated that deficiency of maternal insulin causes an increased influx of mixed nutrients or fuels (glucose, amino acids, lipids, ketones) into fetal circulation resulting in hyperinsulinaemia [4]. Frienkel presented the concept “fuel-mediated teratogenesis” to describe alterations that goes beyond organogenesis causing long-range effects on anthropometric, metabolic and behavioral functions in the offspring due to abnormal fuel mixtures in maternal metabolism due to hyperglycaemia [4]. Studies of developmental origins of health and disease have highlighted the possible role of hyperglycaemic intrauterine environment mediating and accelerating the current epidemic of obesity and diabetes through fetal programming and epigenetic changes [6–9].

While the peripartum and immediate postnatal complications of GDM have been well described, the long-term risks for the offspring have been less studied. Several epidemiologic studies have investigated the association between HIP and offspring anthropometric outcomes during childhood; the majority of them focused on Pima Indians and European and American birth cohorts. Many studies examining the association of offspring BMI with maternal hyperglycaemia in pregnancy have had a small number of exposed offspring thus limiting the power of such studies [10,11]. A large number of studies have reported a positive association between HIP and overweight and obesity [12–27], while few studies have not shown such an association [11,28–30]. Given the limited evidence from South Asian populations for risk estimates for childhood obesity that are attributable to maternal diabetes in utero, further studies in these populations were identified as an important research need [31]. South Asians present with greater metabolic risk at lower levels of BMI compared with other ethnic groups, with type 2 diabetes developing at a younger age and rapidly progressing to other complications [32–34]. Many studies have shown that being obese in childhood and adolescence is associated with obesity in the adult life, and overweight in adolescence is considered an important predictor of long-term morbidity and mortality [28,35–38]. Given the high risk of diabetes and cardiovascular diseases and rising trend of obesity among South Asians, it is imperative that we identify risk groups and target interventions from early life to mitigate the escalating epidemic of non-communicable diseases.

The aim of this study was to determine the association between the intrauterine exposure to hyperglycaemia and anthropometric measurements in offspring at 10 - 11 years of age in Sri Lanka and to determine whether the association was independent of child’s birth weight, parity and mother’s pre-pregnancy BMI.

## Materials and Methods

### Study design and population

A retrospective cohort study was conducted in eight Medical Officer of Health (MOH) areas in Colombo district, Sri Lanka from March 2015 to October 2016 to assess the long term outcomes of HIP on the mother and the offspring. We have previously published the risk of type 2 diabetes in the mothers 10 years after gestational diabetes [39].

Colombo is the most populous district in Sri Lanka with a total estimated population of 2,324,349 amounting to nearly 11% of the total population of the country [40]. For the delivery of public health services, the district is divided into fifteen MOH areas and the metropolitan Colombo Municipal Council area. The total population in the eight MOH areas included in the study was approximately 940,000. Each MOH area is sub divided into Public Health Midwife (PHM) areas, which constitute the smallest field health care delivery unit in the public health system of Sri Lanka. The PHM delivers maternal and child care services as the grass roots level healthcare worker. The PHM maintains a paper-based record keeping system for maternal and child care services and all live births in a given PHM area are recorded in the “Birth and Immunization Register” (BI Register) by the PHM. In the current study, we identified children born in 2005 through the BI registers and through schools in the selected MOH areas.

There was no universal screening programme to screen for HIP in Sri Lanka in 2005. During this period, GDM screening in the antenatal clinics, as per national guidelines at that time, was based on assessment of risk factors (41). These women underwent 75g oral glucose tolerance testing mainly at gestation weeks 24–28. WHO (1999) criteria for 2-hour post 75g oral glucose load (≥140mg/dl) was taken as the criterion for diagnosis of GDM (42).

Since Sri Lanka does not have an electronic database system for keeping patient records and paper-based records are stored only for 5 years in the health institutions, tracing patient held antenatal records to verify exposure status (hyperglycaemia in pregnancy) was the best possible option available. A feasibility study conducted beforehand to verify the availability of patient held antenatal records revealed that approximately 70% of women had antenatal records 10 years after the delivery.

The study was conducted in three stages. In the first stage of the study, a self-administered questionnaire to obtain information on history of hyperglycaemia in the index pregnancy, availability of antenatal records and blood sugar assessment reports of the index pregnancy was sent to all mothers of 2005 born children identified through the BI registers in the community and through schools in the selected MOH areas.

We defined occurrence of hyperglycaemia in the index pregnancy as a positive answer (yes) to the question ‘Did you have high blood sugar / diabetes during the index pregnancy’. Given the high literacy level among women in Sri Lanka, most women were aware of whether they had diabetes during pregnancy.

A total of 7205 children who were born in 2005 were identified in stage 1. The prevalence of self-reported hyperglyceamia in the index pregnancy was 3.5% (N=257). Eighty eight percent (n=226) of mothers of children exposed to HIP still had antenatal records of index pregnancy compared to 69% (n=4811) of mothers of children not exposed to HIP. Potential participants for the main study were identified at the end of the first stage. All children whose mothers had antenatal records and gave a history of HIP during the index pregnancy were considered as “potential participants” to be included in the “exposed group”. For each potential participant in the exposed group, two children of mothers with antenatal records and no history of HIP during the index pregnancy were selected from the same PHM area as “potential participants” to be included in the “non-exposed group”.

During the second stage, the mothers of all potential participants of “exposed” and “non-exposed” groups were invited to participate in the “eligibility assessment sessions”. These eligibility assessment sessions were conducted at PHM area level as it was easily accessible to all mothers thus maximizing participation.

The research team interviewed the mothers of potential participants and scrutinized their antenatal and medical records to identify participants meeting the inclusion criteria (born in 2005, availability of antenatal records, singleton pregnancy) which were previously decided by a group of experts comprising of specialists in obstetrics, obstetric medicine and public health. Having received antenatal care in a unit lead by a Consultant Obstetrician was one of the eligibility criteria for both “exposed” and “non exposed” groups to counter the possibility of misclassification due to limiting the GDM screening to high risk pregnancies in 2005.

170 children exposed to HIP and 291 children not exposed to HIP were identified as eligible and were invited for the study. A sample size of 161 in each group was required to detect a 15% difference in the risk of being overweight with 90% power, an alpha error of 0.05 and a 1:1 ratio between children exposed and not exposed to hyperglcyaemia in utero (23). In the third stage, 159 offspring of women with HIP (OHIP) and 253 offspring of women with no HIP (ONHIP) in the index pregnancy participated in the study. Among the OHIP, 86.8% (n=138) were exposed to gestational diabetes in utero. The detailed flow chart of participant selection is given in Fig 1.

**Fig 1.**
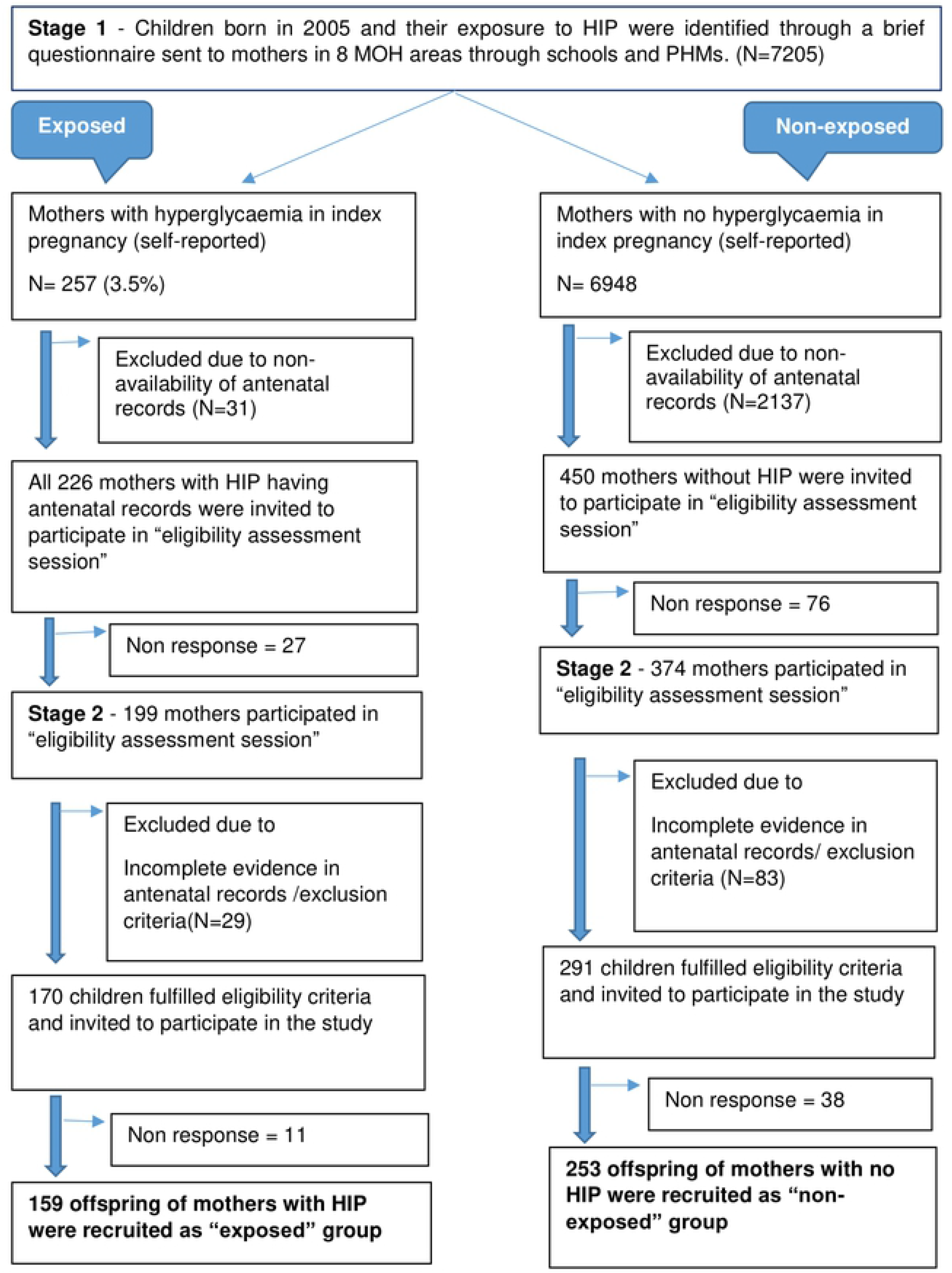
Selection of the study population.

### Data collection

Data collection was carried out by a team of doctors. “Data collection sessions” were arranged in a location easily accessible to participants in a given locality such as a field clinic centres or the MOH office. Socio-demographic characteristics and participants’ physical activity engagement were obtained by interviewing mothers. A 24-hour dietary recall was used to assess the participant’s dietary energy intake. Energy intakes were calculated using the computerized food composition database, FoodBase 2000 software, (Institute of Brain Chemistry, UK) containing Sri Lankan food items and mixed dishes [43] at the Department of Applied Nutrition, Wayamba University of Sri Lanka. Pregnancy related information and glycaemic status during the index pregnancy were extracted from antenatal records to ascertain exposure status using WHO 1999 criteria for diagnosis of diabetes in pregnant women [42].

Anthropometric measurements of the participants were obtained early in the morning following standard operating procedures to ascertain outcome status. Weight and height were measured in light clothing and without shoes. Weight was measured to the nearest 0.1 kg using a calibrated digital scale (SECA 876). Height was measured to the nearest 0.1 cm using a SECA stadiometer. Waist circumference was measured to the nearest 0.1 cm at the mid-point between the lowest rib and the top of the iliac crest with a non-elastic tape. Triceps skinfold thickness was measured to the nearest 0.2mm using a Harpenden skinfold caliper. Two measurements were taken and the mean was used for analysis. The same instruments were calibrated regularly and used throughout the study.

#### Ascertainment of exposure

Children with documentary evidence of exposure to HIP in antenatal records or glucose tolerance tests during the index pregnancy were classified as the OHIP (exposed) group. Diagnosis of GDM and diabetes mellitus in the mother was based on WHO 1999 criteria (42) which was used in Sri Lanka in 2005. Children with no documented evidence of exposure to HIP in antenatal records during the index pregnancy were classified as the ONHIP (non-exposed) group.

#### Ascertainment of outcome

Anthropometric outcome measures were ascertained as follows.

#### Overweight

Overweight was defined as a BMI for age > +1 SD (equivalent to BMI 25kg/m^2^ at 19 years)(44).

#### Obesity

Obesity was defined as a BMI for age > +2 SD (equivalent to BMI 30kg/m^2^ at 19 years) (44).

WHO AnthroPlus for personal computers software for assessing growth of the world’s children and adolescents was used to calculate BMI and BMI z-scores (45,46)

#### Abdominal obesity

Abdominal (central) obesity was defined as waist circumference above the 90^th^ percentile for age and sex (47). Since body fat distribution is different among children of Asian, African and Caucasean races (48), WC percentiles developed for Indian children by Kurian et al (49) were used to identify cutoff values to define abdominal obesity.

#### High Triceps skinfold thickness

High triceps skinfold thickness was defined as TSFT above the 70^th^ percentile for age and sex. Since there are racial differences in skinfold thickness (50), triceps skinfold thickness reference charts developed for Indian children using the same instrument (Harpenden caliper) were used in this study (51).

### Statistical analysis

Baseline characteristics of participants in the OHIP and ONHIP groups were described using descriptive statistics. Variables were tested for normality using the Kolmogorov Smirnov test. Normally distributed continuous data are presented as means (SD) and non-normally distributed data are presented as medians (IQR). Frequencies and percentages were used to summarize categorical variables. Comparisons of baseline and follow up assessment characteristics of OHIP and ONHIP groups were done using t-test (for normally distributed data) or Mann Whitney *U* test (for non-normally distributed data) for continuous variables and the chi square test for categorical variables. Unadjusted Odds ratios and their 95% confidence intervals (CI) were calculated to assess the association between HIP and overweight, obesity, abdominal obesity and high TSFT. Binary logistic regression analysis was carried out to adjust for possible confounding effects of maternal pre-pregnancy BMI, parity and birth weight. All tests of significance were two-tailed. A probability level of P<0.05 was used to indicate statistical significance in all analyses.

### Ethical considerations

The protocol was approved by the Ethics Review Committee of the Faculty of Medicine, University of Kelaniya, Sri Lanka *(Ref. No.P/24/03/2015)*. All mothers of study participants gave informed written consent and verbal assent was obtained from the participants. A “feedback session” was arranged after each data collection session and participants were issued a personal record with anthropometric measurements. All participants and their mothers were counseled on the importance of diet and lifestyle modification for prevention of overweight and cardiovascular diseases. Participants needing specialized care were referred to the Lady Ridgeway Children’s Hospital, a tertiary care facility for children, in Colombo.

## Results

### Characteristics of the study population

A total of 412 children born in 2005 participated in the study. Baseline characteristics of the 159 offspring of women with HIP (OHIP) and 253 offspring of women with no HIP (ONHIP) are compared in table1.

**Table 1.** Characteristics of the exposed (OHIP) and non-exposed (ONHIP) groups

| Characteristic | OHIP group<br>(N=159) | ONHIP group<br>(N=253) | p value |
| --- | --- | --- | --- |
| **Sociodemographic characteristics** |  |  |  |
| Age years - mean (SD) | 10.89 (0.32) | 10.82 (0.31) | 0.009<sup>a</sup> |
| Sex - Male | N=67 (42.1%) | N=118 (46.6%) | 0.37<sup>b</sup> |
| Ethnicity - Sinhala | N=153 (96.2%) | N=231 (91.3%) | 0.24<sup>b</sup> |
| Education level of mother |  |  |  |
| Primary education | N=5 (3.1%) | N=2 (0.8%) | 0.10<sup>b</sup> |
| Secondary education | N=144 (90.6%) | N=241 (95.2%) |  |
| Tertiary education and higher | N=10 (6.3%) | N=10 (4.0%) |  |
| Family Income per month |  |  |  |
| <Rs, 50000 (< USD 280) | N=113 (71.1%) | N=190 (75.1%) | 0.37<sup>b</sup> |
| **Index pregnancy related characteristics** |  |  |  |
| Mother's age at delivery in years – Mean (SD) | 31.9 (5.3) | 27.8 (5.3) | < 0.001<sup>a</sup> |
| Primi Parity | N=53 (33.3%) | N=128 (50.6%) | 0.002<sup>b</sup> |
| Mother's BMI at first trimester of index pregnancy<sup>1</sup> |  |  |  |
| < 18.5 | N=5 (5%) | N= 43 (22.6%) | < 0.001<sup>b</sup> |
| 18.5 – 24.9 | N=56 (55.4%) | N= 131 (69.0%) |  |
| ≥ 25 | N=40 (39.6%) | N= 16 (8.4%) |  |
| Gestational age in weeks – Median (IQR) | 38 (37-39) | 40 (38-40) | <0.001<sup>c</sup> |
| Gestational age at delivery ≥ 37 weeks | N=145 (91.2%) | N=246 (97.2%) | 0.007<sup>b</sup> |
| Birth weight of index child in kg – Mean (SD) | 3.1 (0.5) | 2.9 (0.4) | <0.001<sup>a</sup> |
| Birth weight of index child ≥ 3.5kg | N=39 (24.5%) | N=20 (7.9%) | < 0.001<sup>b</sup> |
| Exclusive breast-feeding duration ≥ 4 months | N= 141 (88.7%) | N= 236 (93.2%) | 0.10<sup>b</sup> |
| **Lifestyle related characteristics** |  |  |  |
| Physical activity (> 1 hour/d for ≥ 5 days/week)<sup>2</sup> | N=71 (44.7%) | N=130 (51.6%) | 0.17<sup>b</sup> |
| Dietary energy intake in kCal – Median (IQR)<sup>3</sup> | 1449 (1238-1864) | 1514 (1257-1850) | 0.78<sup>c</sup> |

<sup>1</sup> Data were available in 63.5% (N=101) of OHIP and 75.1% (N=190) of ONHIP.
<sup>2</sup> Data available for 159 OHIP and 252 ONHIP.
<sup>3</sup> Data available for 70.4% (N=112) OHIP and 74.3% (N=188) ONHIP.
<sup>a</sup> Independent samples T test
<sup>b</sup> Chi square test
<sup>c</sup> Mann-Whitney U test

At the time of the outcome assessment, the age of all participants ranged between 10.3 years to 11.6 years with a mean of 10.85years (SD=0.39). Mothers of children exposed to HIP were older and had significantly higher BMI at the booking visit in the first trimester compared to mothers of non-exposed children (p<0.001). Exposed children were heavier at birth and had a shorter gestational age compared to non-exposed children (p<0.001). About half of the children in ONHIP group were firstborns compared to only one third of children in the OHIP group (p=0.002). Sociodemographic characteristics, breast feeding practices, dietary energy intake and physical activity level were not significantly different between the two groups.

### Outcome assessment

Table 2 compares participants’ anthropometric measurements and prevalence of outcome measures between OHIP and ONHIP groups.

Anthropometric measurements were assessed for normality using 1-sample Kolmogorov-Smirnov test. Height and BMI for age z-score were normally distributed while weight, BMI, WC and TSFT were not normally distributed.

**Table 2.** Anthropometric assessment at follow up

| Characteristic | OHIP group (N=159) | ONHIP group (N=253) | p value |
| --- | --- | --- | --- |
| Height (cm) – Mean (SD) | 141.5 (6.9) | 140.4 (6.9) | 0.09<sup>a</sup> |
| Weight (kg) - Median (IQR) | 34.1 (28.3–41.6) | 28.8 (25.6-36.5) | P<0.001<sup>b</sup> |
| BMI - Median (IQR) | 16.9 (14.7-20.2) | 15.1 (13.8-17.8) | P< 0.001<sup>b</sup> |
| BMI z score – Mean (SD) | -0.1 (1.6) | -0.9 (1.7) | P< 0.001<sup>a</sup> |
| WC (cm) – Median (IQR) | 61.4 (55.7-69.2) | 56.4 (52.7-64.4) | P <0.001<sup>b</sup> |
| TSFT(mm)<sup>1</sup> - Median (IQR) | 13.3 (9.6-17.2) | 9.9 (7.4-14.1) | P < 0.001<sup>b</sup> |

BMI=Body mass index. WC=Waist circumference. TSFT=Triceps Skinfold Thickness.
<sup>1</sup>Data were available for 152 exposed and 236 non-exposed children.
<sup>a</sup>Independent samples T test <sup>b</sup> Mann-Whitney U test

The mean BMI-for-age z-score of exposed children was significantly higher than that of non-exposed children (P<0.001). Exposed children were significantly heavier and had significantly higher median BMI, WC and TSFT than the non-exposed children (p< 0.001). Fig 2 depicts the distribution of anthropometric parameters.

**Fig 2.**
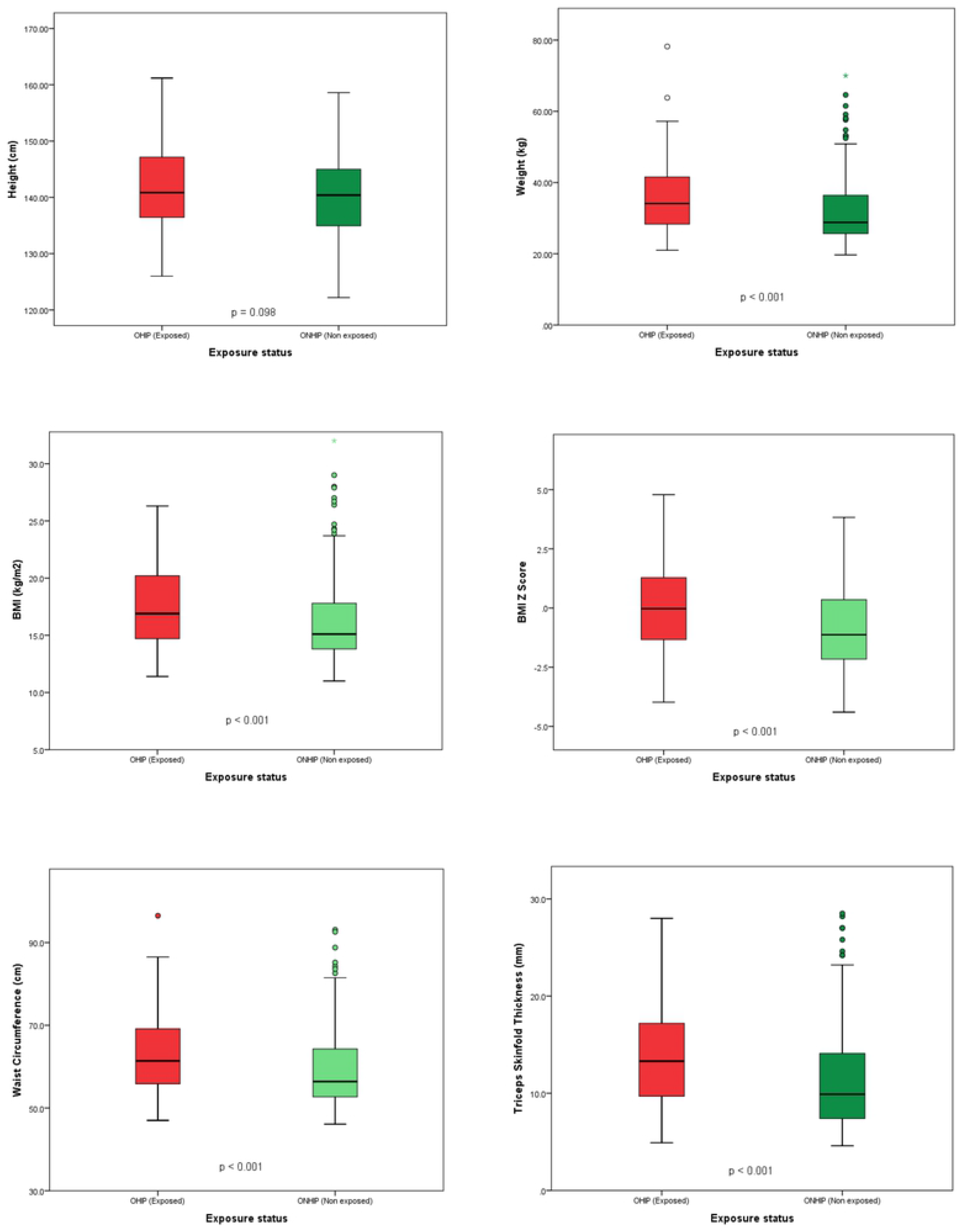
Distribution of anthropometric parameters.

Table 3 shows the anthropometric outcome status of children at follow up.

**Table 3:** Outcome status of participants at follow up

| Outcome status | OHIP group (N=159) | ONHIP group (N=253) | Odds Ratio<br>(95% CI of OR) | p value |
| --- | --- | --- | --- | --- |
|  | Prevalence (95% CI) | Prevalence (95% CI) |  |  |
| Overweight (BMI<sup>a</sup><br>z score > +1SD) | 30.8% (23.6 - 37.9) | 16.2% (11.6 - 20.7) | 2.3(1.4 -3.7) | p<0.001 |
| Obesity (BMI<sup>a</sup> z score<br>> +2SD) | 5.7% (3.0-10.4) | 5.1% (3.0-8.6) | 1.1 (0.4 -2.6) | P=0.82 |
| Abdominal obesity<br>(WC<sup>b</sup> ≥ 90th percentile) | 15.1% (9.5 – 20.6) | 7.1% (3.9 – 10.2) | 2.3 (1.2 -4.4) | p=0.009 |
| TSFT<sup>c</sup>>70<sup>th</sup> percentile | 36.2% (28.5 – 43.8) | 20.8% (15.6 – 25.9) | 2.2 (1.4– 3.4) | p=0.001 |

<sup>a</sup>BMI= Body mass index
<sup>b</sup>WC= Waist circumference
<sup>c</sup>TSFT=Triceps skinfold thickness.

The prevalence of overweight, abdominal obesity and high TSFT were significantly higher among the offspring of mothers who had HIP. The high prevalence of abdominal obesity (7.1%) and high TSFT (20.8%) even among the children not exposed to HIP is a concern. Children exposed to HIP were 2 times more likely to be overweight and have abdominal obesity and have a TSFT > 70^th^ percentile than non-exposed children (p < 0.01). Prevalence of obesity was similar in both groups.

### Association between HIP and anthropometric outcome measures after adjusting for confounders

Logistic regression analysis was carried out to describe the association between the HIP and anthropometric outcome status (overweight, abdominal obesity, TSFT >70^th^ percentile) in 10-11year old children after adjusting for maternal BMI in the first trimester, parity of index pregnancy and birth weight. Predictors of overweight, abdominal obesity and high TSFT in the offspring are given in Table 4.

**Table 4:** Predictors of anthropometric outcome status.

| Risk factor | Overweight<br>(BMI z score > +1SD) |  | Abdominal obesity<br>(Waist circumference ≥<br>90th percentile) |  | TSFT > 70<sup>th</sup> percentile |  |
| --- | --- | --- | --- | --- | --- | --- |
|  | Unadjusted | Adjusted | Unadjusted | Adjusted | Unadjusted | Adjusted |
|  | OR<br>(95% CI) | OR †<br>(95% CI) | OR<br>(95% CI) | OR †<br>(95% CI) | OR<br>(95% CI) | OR †<br>(95% CI) |
| Exposure to HIP | 2.3 **<br>(1.4-3.7) | 2.5*<br>(1.3-4.7) | 2.3*<br>(1.2-4.4) | 2.9*<br>(1.2-6.8) | 2.2**<br>(1.4-3.4) | 2.1*<br>(1.2-3.9) |
| Maternal BMI ≥<br>25kgm<sup>2</sup> in the first<br>trimester | 3.3**<br>(1.8– 6.2) | 2.8*<br>(1.4-5.8) | 2.4*<br>(1.1– 5.6) | 1.9<br>(0.7-4.7) | 2.9**<br>(1.6– 5.4) | 2.4*<br>(1.2-4.8) |
| Firstborn child | 1.6*<br>(1.01-2.6) | 2.6*<br>(1.4-4.9) | 1.4<br>(0.7-2.7) | 1.8<br>(0.8-4.1) | 1.3<br>(0.8-2.1) | 2.1*<br>(1.2-3.8) |
| Birth weight ≥ 3.5kg | 1.9*<br>(1.03-3.4) | 1.6<br>(0.7-3.5) | 0.7<br>(0.2-2.1) | 0.6<br>(0.2-2.1) | 2.2*<br>(1.2-3.9) | 2.6*<br>(1.3-5.4) |

HIP=Hyperglycaemia in pregnancy TSFT=Triceps skinfold thickness BMI=Body mass index
† Adjusted for maternal BMI, parity, birth weight and exposure to HIP.
\*P < 0.05, \*\*P < 0.001

Even after adjustment for maternal BMI, birth weight and birth order, exposure to HIP was a significant predictor of overweight, abdominal obesity and high TSFT in the offspring at 10 years of age. Maternal overweight in the first trimester, a proxy for pre-pregnancy overweight, is an independent risk factor for offspring overweight and high TSFT at 10-11 years. Similarly, being the first-born child carries a more than two-fold increased risk of overweight and high TSFT independent of maternal BMI, birth weight and exposure to HIP.

## Discussion

To the best of our knowledge, this is the first study on long term implications of HIP on anthropometric parameters in the offspring in Sri Lanka and one among the handful of studies from South Asia. Even the previous studies conducted in India (26,52) were limited by the small number of offspring of GDM mothers (n=41 and n=35). The significant associations between maternal HIP and overweight, abdominal obesity and high TSFT in the offspring in this study support the hypothesis that intrauterine exposure to HIP may have a long term risk of increased adiposity in the offspring. The higher BMI and BMI-z-score in the offspring of women with HIP reported in this study is consistent with earlier studies (14,16,21,22,53–56). A comprehensive meta-analysis by Philipps et al. identified a strong association between intra uterine exposure to maternal diabetes and increased offspring BMI in childhood (10). The prevalence of overweight (BMI-z-score > +1SD) was significantly higher among OHIP compared to ONHIP (30.8% vs 16.2%). Our results are similar to findings of other studies that have reported a higher risk of overweight and obesity among offspring of mothers who had HIP (11,14,17,18,21–25,57–60). However, in contrast to other studies, the prevalence of obesity (BMI-z-score > +2SD) was similar in the exposed and non-exposed groups in our study.

In our study, children exposed to intrauterine hyperglycaemia had a significantly higher waist circumference at 10 years compared to non-exposed children. Previous studies have reported similar findings of significantly higher waist circumference among offspring exposed to hyperglycaemia in utero including a multinational study involving 206 offspring of GDM mothers and 4534 offspring of non-GDM mothers from 12 countries (24,61,62).

In our study, children exposed to HIP had significantly higher TSFT than children not exposed to HIP (13.3mm vs 9.9 mm; p< 0.001). Wright et al, observed that children exposed to GDM had significantly higher sum of skinfold thicknesses (Subscapular and Triceps) than non-exposed children (63). Cumme et al, reported increased subscapular to triceps skinfold thickness ratio in children exposed to HIP (62). Krishnaveni et al. from India, observed significantly higher TSFT among the offspring of diabetic mothers compared to offspring of non-diabetic mothers at 5 years of age (26). When the same cohort was assessed at 9.5 years of age, they observed a significantly higher BMI and TSFT among girls exposed to intrauterine hyperglycaemia but not among boys (14). No significant difference between the growth of the boys and girls was observed in our study (results not shown).

In contrast to the many studies where the association between maternal HIP and child overweight attenuated towards the null after adjusting for maternal BMI (11,56,57)(62), our results were statistically significant even after adjusting for maternal BMI, child’s birth weight and birth order.

We included offspring of women with any type of HIP (gestational diabetes, pre-existing diabetes or overt diabetes first detected in pregnancy) in the “exposed” group without stratification by type of diabetes based on previous research which showed that long-term consequences of HIP on offspring overweight are independent of mother’s diabetes type (25,64,65). A sub-group analysis of a meta-analysis by Philips et al, revealed that there is no difference in offspring BMI-z-score in relation to diabetes type such as GDM or Type 1 diabetes(10).

Using three methods (BMI, waist circumference and triceps skinfold thickness) to assess adiposity of participants is a unique strength of this study. BMI is widely used to measure body composition and is used to define overweight and obesity (66). Though BMI is widely used to measure generalized obesity or adiposity, its value in discriminating lean body mass from fat mass has been challenged (67). Skinfold thickness is a valid measurement of subcutaneous fat (68) and there is evidence to suggest that later adulthood adiposity is better predicted by adolescent skinfold thickness than by adolescent BMI (69). In predicting cardiovascular disease risk, abdominal adiposity appears to be superior to BMI (66). Abdominal obesity, defined as waist circumference >90^th^ percentile is a mandatory criterion for diagnosing metabolic syndrome in children and adolescents (70). Since body fat distribution is different among children of Asian, African and Caucasean races (48), waist circumference percentiles developed for Indian children based on measurements made on 9060 children 3-16 years of age (49) were used to identify cutoff values to define abdominal obesity in the present study. Since there are racial differences in skinfold thickness (50), triceps skinfold thickness reference charts developed for Indian children using the same instrument (Harpenden caliper) were used in this study (51).We decided to use the TSFT >70^th^ percentile as the cut off for “high triceps skinfold thickness” based on the findings of the same study where they identified the 70^th^ percentile as the cutoff for predicting risk of hypertension in children (51).

Having a large number of offspring exposed to HIP is a major strength of our study. Selecting both “exposed” and “non-exposed” children from the same source population in the community based on antenatal records reduced recall bias and misclassification. Since exposure was assigned on an earlier date than the outcome was measured in the child, it is unlikely that the outcomes of interest would have influenced the classification of exposure status. Children whose mothers received antenatal care from a consultant obstetrician were selected in both exposed and non-exposed groups. Since universal screening for HIP was not available in 2005, having being under the care of a Consultant obstetrician implies that they had a fair chance of being screened and diagnosed for HIP, if required, thus minimizing misclassification bias. Even if misclassification did occur, the associations between HIP and anthropometric outcome measures we observed is likely to be an underestimation.

Not having detailed information on maternal blood sugar levels at diagnosis and glycaemic control during pregnancy is a limitation of this study. In general, all women diagnosed to have HIP are advised on dietary management and physical exercise. Those women who cannot obtain satisfactory glycaemic control with lifestyle management alone are started on pharmacological management with metformin or insulin. For the purpose of this study, we have collected data on whether the mothers were on diet control alone, started on metformin or on insulin from antenatal records.

Missing maternal pre-pregnancy BMI data on nearly 30% of mothers is another limitation. Maternal BMI in the first trimester was used as a proxy for pre-pregnancy BMI. According to national maternal care guidelines in Sri Lanka (41), BMI is measured and recorded as three categories (<18.5, 18.5 – 24.9, ≥ 25) in the first clinic visit only if the woman presents before the completion of the 12^th^ week of gestation. It is likely that some of these women whose BMI data were not available would have presented for the booking visit after 12 weeks of gestation. As the data were extracted from the antenatal records, we had to limit to the above 3 categories of BMI when adjusting for maternal BMI. It would have been ideal if we adjusted for the weight gain in pregnancy. But this data was not available for the majority of the participants. We adjusted for the birth weight of the child which can be taken as a proxy measure for weight gain in pregnancy. Based on the national guidelines on antenatal care in Sri Lanka, birth weight ≥ 3.5kg was taken as macrosomia (41).

The results of this study have several important public health implications. Locally generated evidence in this study would be an eye opener for clinicians, field health care workers and health policy makers to take necessary actions to follow up exposed children closely during the critical period of development to prevent and to detect the appearance of anthropometric risk parameters early. Creating awareness on possible long term effects of maternal hyperglycaemia would motivate women to achieve better glycaemic control during pregnancy and lifestyle modification of the child with adherence to a healthy diet and increased physical activity to reduce the risk of overweight. Given the high prevalence of HIP in Sri Lanka and other South Asian countries, preventive strategies targeted at women of childbearing age and offspring of women with HIP are likely to have a significant population health impact on the current epidemic of obesity and non-communicable diseases.

## Conclusions

Children exposed to intrauterine hyperglycaemia have higher BMI, waist circumference and TSFT at 10-11 years compared to children who were not exposed independent of maternal pre-pregnancy overweight, birth weight and birth order. It is imperative to implement long term follow up for children exposed to hyperglycaemia in pregnancy with anthropometric assessment and life style modification advice to reduce the risk of developing overweight and associated metabolic and cardiovascular disturbances.

## Acknowledgements

Authors would like to acknowledge the National Research Council of Sri Lanka for providing the research grant and the team of doctors (Dr Chathura Edirisinghe, Dr Kojika Withana, Dr Chamini Sumanasena, Dr Sachithra Dilrukshi and Dr Chamidu Wickramathunga) for their support as research assistants during data collection.

## Supporting information

S1 Dataset. HIP and risk of adiposity in the offspring at 10 years - Sri Lanka

